# Developing oak buds produce volatile emissions in response to herbivory by freshly hatched caterpillars

**DOI:** 10.1101/2023.10.18.562884

**Authors:** Jessica L. Graham, Michael Staudt, Bruno Buatois, Samuel P. Caro

## Abstract

Plant responses to damage by insectivorous herbivores is well documented in mature leaves. The resulting herbivore-induced plant volatiles (HIPVs) protect the plant by attracting carnivorous arthropods and even some insectivorous vertebrates, to parasitize or consume the plant invaders. However, very little is known about plant production of HIPVs in developing buds, particularly when herbivorous insects are too small to be considered a prey item. It is additionally unclear whether plants respond differently to generalist and specialist chewing insects that overlap in distribution. Therefore, we compared HIPV production of Downy oak (*Quercus pubescens* Willd.) buds infested with freshly hatched caterpillars of *Tortrix viridana* (specialist) and *Operophtera brumata* (generalist), against uninfested buds. Of the compounds identified in both years of the experiment, we found that (*Z*)-hex-3-enyl acetate, (*E*)-β-ocimene, acetophenone, linalool, (*E*)-4,8-dimethyl-1,3,7-nonatriene (DMNT), methyl salicylate, *α*-copaene, *α*-humulene, (*E*)-caryophyllene, and (*E,E*)-*α*-farnesene were higher in infested buds compared to controls. We found no difference in HIPV production between the specialist and the generalist herbivores. Production of HIPVs was also associated with leaf damage, with higher HIPV production in more severely attacked buds. Thus, our study shows that oak trees already start responding to insect herbivory before leaves are developed, by producing compounds similar to those found in damaged mature leaves. Future work should focus on how Downy oak may benefit from initiating alarm cues at a time when carnivorous arthropods and insectivorous vertebrates are unable to use herbivorous insects as host or food.

## Introduction

Plants have developed multiple methods to reduce damage caused by herbivory. One of these methods is through the release of herbivore-induced plant volatiles (HIPVs). These airborne signals have been shown to attract carnivorous arthropods (Turlings & Benrey 1998; Van Poecke *et al*. 2001) and even some insectivorous vertebrates (Amo *et al*. 2013; Mäntylä *et al*. 2017; Goldberg *et al*. 2019). Using these chemical signals, carnivorous arthropods and insectivorous vertebrates can significantly reduce the number of herbivorous insects, thus preventing further damage to the plant (Kessler & Baldwin 2001).

While a wealth of studies have described herbivore-induced volatile emissions from mature foliage, only a few have specifically investigated the emissions from infested buds or very young developing shoots (Röse & Tumlinson 2004; Brilli *et al*. 2009, Rostas & Eggert 2008). Generally, buds are well protected against biotic and abiotic stresses during the dormancy period. The tiny embryo shoots they contain are densely enveloped by tough scales impregnated with gummy or resin type substances, efficiently impeding attacks by insects or pathogens. However, buds become vulnerable during bud break when the scales are spreading or fall off giving access to the inner soft tissues of the developing shoot. Herbivore attacks on these organs can be very detrimental to the plants (Alves-Silva & Del-Claro 2016), because even small damages will cause the complete loss of a new shoot for the coming growing season and not only the fraction of an already existing leaf (Blundell & Peart 2000; Kozlov & Zvereva 2014). Bud herbivory and parasitism particularly threaten deciduous woody plants growing in climates having a very short vegetation period, which limits both the replacement of lost shoots by re-sprouts from dormant buds and the acquisition of resources by the remaining foliage. For example, in the Mediterranean area, the growth of new shoots is restricted to the period of spring due the onset of drought in summer (Montserrat-Marti *et al*. 2009).

Although damages inflicted by insect herbivores to buds are presumably more detrimental to the plant than damages inflicted to mature leaves, it is unclear whether HIPV emission is adaptive at this early stage. Buds are mostly present in early spring, at a time when many herbivorous insects like caterpillars only start to hatch from eggs and are therefore still small (van Asch & Visser 2007). At that time, those herbivores might not be valuable enough for predators to consider them as prey items. Considering that the fitness benefits for plants in releasing HIPVs are not always evident, even for mature leaves (Cipollini & Heil 2010; Dicke & Baldwin 2010), the net benefit of emitting HIPVs at the bud stage may be even more questionable. Therefore, to better understand the potential benefits provided by HIPV production, we must first clarify whether developing buds respond to herbivory by releasing HIPVs (Röse & Tumlinson 2004; Brilli *et al*. 2009).

Many herbivorous insects are specialized to feed their larvae on the developing buds and foliage of their host trees (Cízek 2005). Thus, plants are unlikely to be exposed to a single herbivorous insect. For example, the green oak tortrix (*Tortrix viridana*) and the winter moth (*Operophtera brumata*) are serious defoliators of European deciduous tree species (Schroeder & Degen 2008) and specifically attack developing buds and shoots in early spring (Du Merle 1999, Hunter 1990). While the green oak tortrix is a *Quercus* specialist, limited to oak stands (Du Merle 1999), winter moths are polyphagous and capable of consuming up to 100 different plant species (Hunter 1990). It is suggested that plants are able to fine-tune the attractiveness of their emissions to carnivores (Dicke & Baldwin 2010) by adjusting their emissions to the type or level of damages they experience (Alborn *et al*. 1997; De Moraes *et al*. 1998; Gaquerel *et al*. 2009; Bricchi *et al*. 2010; Girling *et al*. 2011; Zebelo *et al*. 2014). Whether plants differ in HIPV emissions among chewing herbivorous insects has not been extensively studied yet (e.g. De Moraes *et al*. 1998). With newly emerging evidence that polyphagous caterpillar species like *Helicoverpa zea* produce a salivary enzyme that inhibits the emission of HIPVs in host plants (Lin *et al*. 2021), there may be differences in HIPV responses between specialist herbivores that co-evolve with their host, and polyphagous herbivores with a broad diet.

In the present study, we investigated whether plants release HIPVs in response to insect herbivory during the early stages of leaf development by infesting developing *Quercus pubescens* oak buds with freshly hatched caterpillars and comparing the HIPV emissions of caterpillar infested and uninfested buds. Additionally, we examined if HIPVs released by oak buds early in the season differ between chewing herbivore species by comparing an oak specialist (green oak tortrix) and polyphagous (winter moth) species, and we investigated whether HIPV emissions are proportional to the damages herbivores inflict to the buds.

## Methods

### Plants

During the spring of 2017 and 2018, 80 Downy oak trees (*Quercus pubescens* Willd.) grown in pots were maintained on the north side of a building (*n* = 20), south side of a building (*n* = 20), or at a higher elevation to delay bud burst (*n* = 40). All trees were treated with copper in a Bordeaux mixture (AMM 9500302) and a rapeseed oil insecticide (Naturen Eradibug, AMM 2110150) in the winter preceding each experiment to reduce the chance any fungus and wild insects infested the trees and caused other volatile emissions than the ones induced by our caterpillar infestation.

Bud stages were defined using a slightly modified classification from Du Merle & Mazet (1983). Briefly, bud stages ranged from 1 to 9, with 1 corresponding to a winter dormant bud, and 9 to a fully developed leaf. Stage 4 corresponds to a bud that elongates and loses its scales, and stage 5 corresponds to a bud that opens up. Once at stage 4, the oak buds were considered ready to use in the experiment as caterpillars are unable to perforate the protective scales to enter the bud prior to that stage.

### Insects

Winter moth (*Operophtera brumata*) and green oak tortrix (*Tortrix viridana*) adult females lay their eggs next to leaf buds, which hatch in spring contemporary with bud break (Du Merle 1999). Winter moth is a more generalist herbivore adapted to cooler temperate climates and is considered as an invasive species in North America, while green oak tortrix is rather adapted to warm temperate climates and feeds mainly on few species overall oaks. The green oak tortrix is widely distributed throughout western Europe, though as a *Quercus* specialist, limited to oak stands (Du Merle 1999). Winter moths are polyphagous, but compete with green oak tortrix for oak hosts in parts of their distribution (Hunter 1990). Green oak tortrix and winter moth eggs were collected the previous summer from Corsica, France and Gelderland, The Netherlands, respectively. Eggs were kept in an unheated building overwinter and moved outdoors in spring, so caterpillars and trees were experiencing the same weather conditions.

### Volatile Sampling

Push-pull dynamic headspace sampling was used to collect biogenic volatile organic compound (VOC) emissions from infested and uninfested (control) oak buds (Tholl *et al*. 2006). A double-walled glass headspace (170 cm^3^) enclosed the twig with the focal bud(s). The headspace was closed with a silicone cap pierced with three holes, one holding the twig, and two holding PFA ¼” tubes allowing air entrance and exit from the headspace. Before entering the headspace, ambient air was pumped through a charcoal filter to remove any contaminants. A flow meter (VAF-G2-03M-1-0, Swagelock, Lyon Vannes et Raccords, Genas, France) was used to control the flow of air into the headspace (260 mL min^-1^). Water from a thermo-stated water bath was circulated through the outside of the headspace to maintain a temperature of 25°C inside the headspace as temperature strongly affects volatile emissions from plants (Niinemets *et al*. 2011; Peñuelas & Staudt 2010; Vallat *et al*. 2005).

One day before sampling, the focal trees were moved to the system to allow any volatile emissions produced while moving to dissipate. When the system was initially turned on, clean air was pumped freely through the system for 10 minutes to purge the headspace. The terminal part of an intact twig (holding 1-8 buds, see below) was then placed carefully into the headspace. The system continued to run for at least 30 min to clear any stress-induced compounds released by the plant when placing the buds into the headspace and allow volatile emissions to come to a steady state. A Gilian GilAir air sampler (Sensidyne, LP, St. Petersburg, FL, USA) was attached to the system and pulled air from the headspace at 200 mL min^-1^ into a thermal-desorption stainless-steel cartridge (Perkin Elmer, Villebon-sur-Yvette, France) containing 280 mg of Tenax^®^ TA adsorbent (20/35 mesh, Alltech-Grace Davison, Düren, Germany) to collect volatile emissions. The net influx of clean air kept ambient air from entering the system. Sampling lasted for 2h 30min to collect volatiles from ∼30L of air. In addition to sampling buds (with or without caterpillars), a blank sample was collected each day to subtract background compounds from the bud samples collected that day. These samples were collected in the same way as the bud samples, just without anything in the headspace. Cartridges were stored at -20°C until analysis.

Thirty-three oak trees were sampled in 2017 (*n* = 14 *tortrix* infested, *n* = 10 winter moth infested, *n* = 9 control) and 2018 (*n* = 10 *tortrix* infested, *n* = 12 winter moth infested, *n* = 11 control). A healthy, well-developed terminal twig with 1 – 8 buds was selected for analysis on each tree (*n* = 42 with 1 bud, *n* = 11 with 2 buds, *n* = 4 with 3 buds, *n* = 5 with 4 buds, *n* = 1 with 5 buds, *n* = 2 with 6 buds, *n* = 1 with 8 buds). Each twig was repeatedly measured before (T_0_) and after caterpillar infestation at 24 h (T_24_) and 48 h (T_48_) in 2017, and 48 h (T_48_) and 96 h (T_96_) in 2018. Bud stage for first measurement ranged between 4 and 7 in both years. After the first measurement, the focal buds were either infested with green oak tortrix caterpillars, winter moth caterpillars, or left as a control. Control twigs were thus repeatedly sampled the exact same way, except that they were never infested with caterpillars.

A paintbrush was used to gently place newly hatched caterpillars on the buds where they remained until the end of sampling. Though caterpillars were observed entering the buds in 2017, several buds had no caterpillars remaining at the end of sampling (*n* = 2 *tortrix* infested, *n* = 3 winter moth infested). Thus, in 2018, wire cages with mesh cloth bags were attached to the branch with the focal bud to prevent caterpillars from dispersing. Cages were attached the same way to the sampled branches of control trees.

### Gas Chromatography Mass Spectrometry (GC-MS)

Analyses were carried out at the PACE (Plateforme d’Analyses Chimiques en Ecologie), with the support of LabEx CeMEB, an ANR “Investissements d’avenir” program (ANR-10-LABX-04-01). Cartridges were placed on a TD-20 two-stage thermal desorption system coupled to a QP-2010-SE gas chromatography mass spectrometry system (GC-MS; Shimadzu Corporation, Kyoto, Japan). VOCs were first thermally desorbed by flushing the cartridge heated at 250°C during 10 minutes with a 30mL min^-1^ flow of helium, and absorbed for the second stage on a - 10°C low-dead volume cold trap containing Tenax^®^ TA adsorbent (80/100 mesh, Shimadzu Corporation, Kyoto, Japan). The cold trap was subsequently heated to 250°C for 5 minutes. VOCs were then injected into a 250°C heated transfer line in the GC-MS column (Optima-5MS, 30 m x 0.25 mm x 0.25 µm, Macherey-Nagel, Düren, Germany). The oven temperature program was as follow: 40°C held for 2 minutes, then increased at a rate of 5°C min^-1^ until reaching 200°C, and finally increased at a rate of 10°C min^-1^ up to 270°C held for 6 minutes. Helium was used as carrier gas at 36.1cm s^-1^. The temperature of the transfer line and ion source were 250 and 200°C, respectively. MS was then used with electron impact at 70eV in the scan mode, from 35 to 350 m/z.

Peaks were identified by comparing their retention indices calculated with alkanes (Sigma Aldrich®, Darmstadt, Germany) and mass spectra with those of commercial databases (Adams, 2007 and NIST 2005, respectively). Pure standards dissolved in methanol were used to i) confirm identification and ii) make three-level calibration curves of four specific standards, representing classes of VOCs, in order to estimate emissions (*α*-pinene, *(E)*-caryophyllene, methyl-salicylate and (*Z*)-hex-3-enyl acetate for monoterpenoïds, sesquiterpenoïds, shikimic acid and fatty acids derivatives, respectively). The class mean calibration factor (ng unity^-1^ of peak area) were applied to quantify the peaks identified in the corresponding classes, and the mean calibration factor of all the standards was applied to peaks to unidentified peaks.

Area under the curve (ng) divided by volume of air sampled (L) was used to calculate the concentration for each compound (ng.L^-1^). Since the air flow in the headspace was always kept constant, changes in the net HIPV concentrations must reflect changes in the emission rate per enclosed shoot.

### Bud Damage

After sampling was complete in 2018, infested buds (*n =* 21) were dissected to assess the level of damage to the bud after being infested for 96 h. Buds were dissected and damaged leaves were photographed against graph paper and analyzed using ImageJ software (Schneider *et al*. 2012). Leaf damage area (mm^2^) was measured for all 21 samples from infested trees. Buds sampled later in the season (*n =* 13) have photographs of both the damaged and undamaged leaves that allowed us to calculate damaged leaf area divided by total leaf area (i.e., proportion of total bud damage). However, using this measure reduced sample size. We found a significant correlation between damaged area (mm^2^) and proportion of total bud damage in the 13 samples with both measures (F_1,11_ = 35.48, *p* < 0.001, r^2^ = 0.76). Thus, damaged area (mm^2^) is used in analyses reported here.

### Statistical Analyses

All analyses were conducted in R version 3.5.1 (R Core Team 2020). Initially, 68 and 52 volatile organic compounds were found in at least 1 sample in 2017 and 2018, respectively. We reduced our analyses to the following compounds that were identified in both years: (*Z*)-hex-3-enyl acetate, (*E*)-β-ocimene, acetophenone, linalool, (*E*)-4,8-dimethyl-1,3,7-nonatriene (DMNT), methyl salicylate, *α*-copaene, (*E*)-caryophyllene, (*E,E*)-*α*-farnesene, (*Z)*-hex-3-enyl benzoate, (*E)*-nerolidol, and dendrolasin. In addition, we included aromadendrene and *α*-humulene that were only regularly found in 2017 samples, but that have been associated with larval feeding (Martins & Zarbin 2013; Pinto-Zevallos *et al*. 2013). Two samples contained values greater than nine standard deviations from the mean and were thus, excluded from analyses.

To identify the experimentally controlled factors influencing VOC variation, data was analyzed using redundancy analysis (RDA) (Hervé *et al*. 2018). Prior to this analysis, a small constant was added to the data to remove zeroes. Data was pre-transformed using centered log ratio (package: Hotelling, Curran 2018) and then auto-scaled (Hervé *et al*. 2018). An RDA model was fit with treatment (winter moth infested, tortrix infested, or control), time (T_0_, T_24_, and T_48_), and the interaction of treatment and time using the package vegan (Oksanen *et al*. 2019). We excluded T_96_ samples, which were only collected in 2018, from the analysis because the non-detection of aromadendrene and *α*-humulene in 2018 required entering missing values as zeros. We did not want to include a sampling time that would increase the likelihood of type I error for these compounds by having a mean of 0 ng/L at T_96_ compared to positive, non-zero values in all other sampling times. The condition function was used to control for repeated measures in individual trees and across years.

Using the package RVAideMemoire (Hervé 2019), a constrained and unconstrained principle component analysis (PCA) was run on the RDA model. The constrained PCA determined the proportion of the variance that was explained by the controlled variables in the experimental design (treatment, time, and the interaction of treatment and time) and the unconstrained PCA to determine the proportion of the variance explained by unknown factors that were not included as controlled variables. An analysis of variance (ANOVA) *F*-test with 999 permutations tested whether the proportion of variance explained by the controlled variables was significant. In the case of significant models, a multivariate ANOVA was used to determine which controlled variables from the model (treatment, time, and the interaction between treatment and time) contributed significantly to the proportion of variance explained by the experimental design. Pairwise tests were conducted on significant variables with more than two factors to determine which groups were significantly different from each other. In groups that were significantly different, a biplot was produced to visualize which compounds appeared to be driving the observed differences.

When analyzing leaf area damage, all analyses were restricted to the 96hr sampling point from infested trees only (*n* = 21 samples). Controls were excluded because we assumed no damage was done to the buds. We first ran a linear model to determine if one species of caterpillar caused more damage to the buds than the other. To further identify if caterpillar species or damaged leaf area influenced HIPV production, we ran a second linear model to measure the effects of damage area, species, and the interaction of damage area and species on changes in overall HIPV emissions from the plants. For each sample, the emissions for all 12 HIPVs identified in the 2018 samples were summed to avoid any issues arising from repeated analyses and collinearity between HIPV emissions. Mean values are reported ± SEM. We set α = 0.05.

## Results

Table 1 provides details of VOC concentrations from control, green oak tortrix infested, and winter moth infested buds.

**Table 1.**
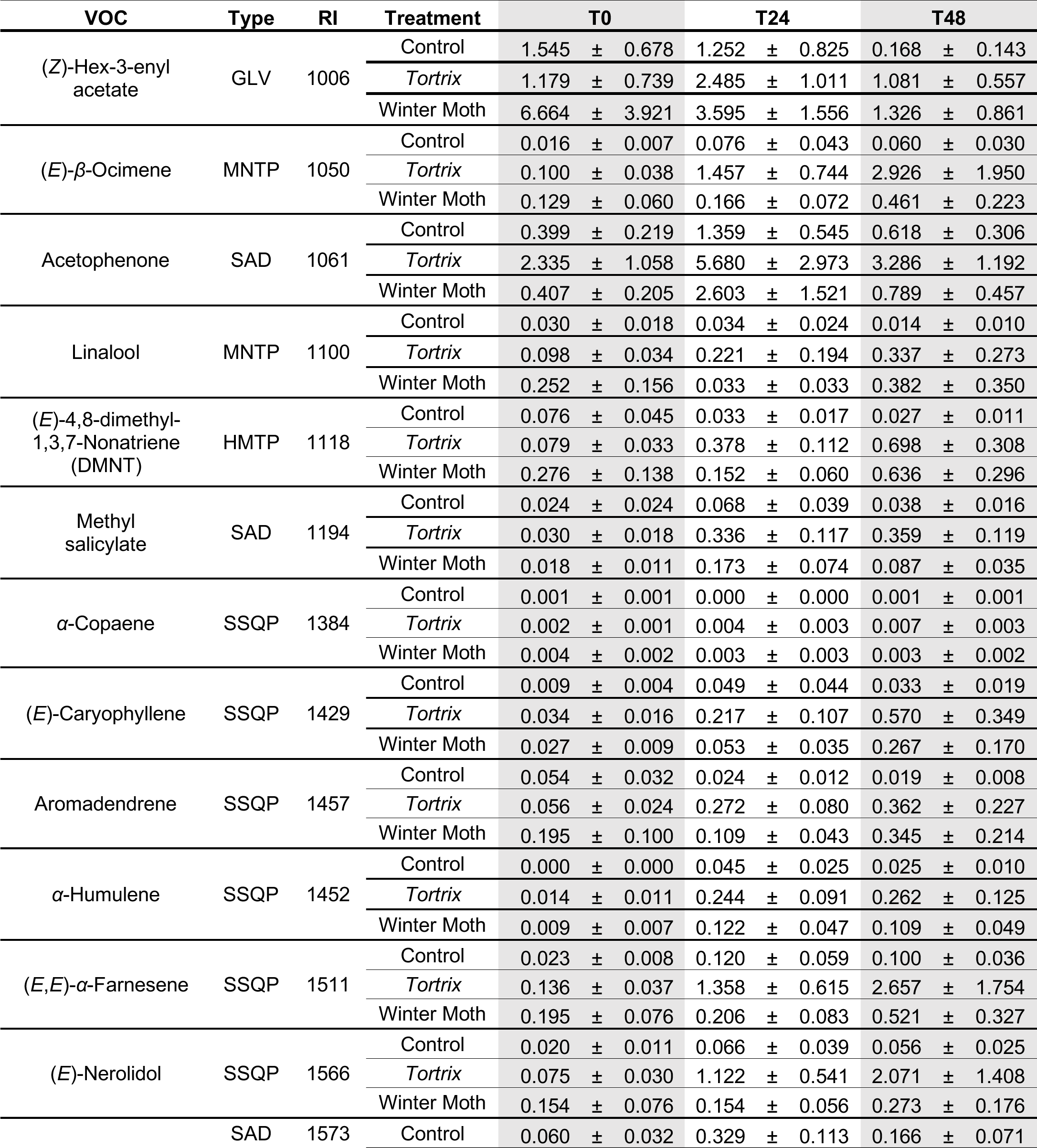
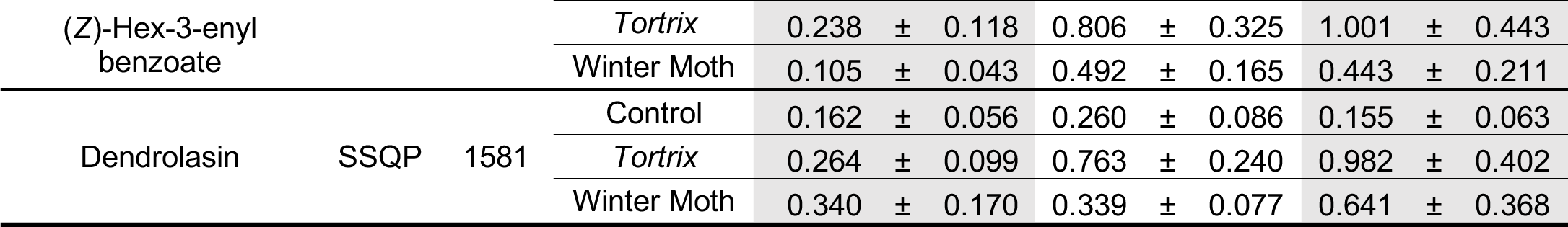
Mean concentrations (ng/L ± SEM) of the 14 HIPVs identified for analysis from the three sampling times for control, green oak tortrix (*Tortrix*) infested, and winter moth infested buds. GC-MS linear retention time (RI) and type of volatile organic compound are also included. GLV = green leaf volatile; HMTP = homoterpene; MNTP = monoterpene; SAD = shikimic acid derivative; SSQP = sesquiterpene.

### HIPV Emissions

The variance explained by the experimental design was significant (constrained variance = 7.81%, *F*_1,8_ = 1.97, *p* = 0.001). Treatment (*F*_1,2_ = 1.84, *p* = 0.014) and time (*F*_1,2_ = 4.06, *p* = 0.001) were both significant, but the interaction between treatment and time was not (*F*_1,4_ = 0.99, *p* = 0.46). Pairwise comparisons of treatment found that the controls differed significantly from both *tortrix* (*p* = 0.015) and winter moths (*p* = 0.015), but *tortrix* and winter moths were not significantly different from each other (*p* = 0.825). Pairwise comparisons of time showed T_0_ significantly differed from T_24_ (*p* = 0.003) and T_48_ (*p* = 0.003). T_24_ significantly differed from T_48_ (*p* = 0.011). While there was overlap between VOCs produced in caterpillar infested and control buds, the following compounds appeared to significantly differ in caterpillar infested buds compared to uninfested buds (figure 1): (*E*)-β-ocimene, linalool, methyl salicylate, *α*-humulene, DMNT, acetophenone, (*E,E*)-α-farnesene, (*E*)-caryophyllene, (*Z*)-hex-3-enyl acetate, and *α*-copaene. Compounds that appear to be borderline significant between controls and caterpillar infested buds were aromadendrene and (*Z*)-hex-3-enyl benzoate. Non-significant VOCs included dendrolasin and (*E*)-nerolidol. Therefore, we consider all VOCs except dendrolasin and (*E*)-nerolidol as HIPVs.

**Figure 1.**
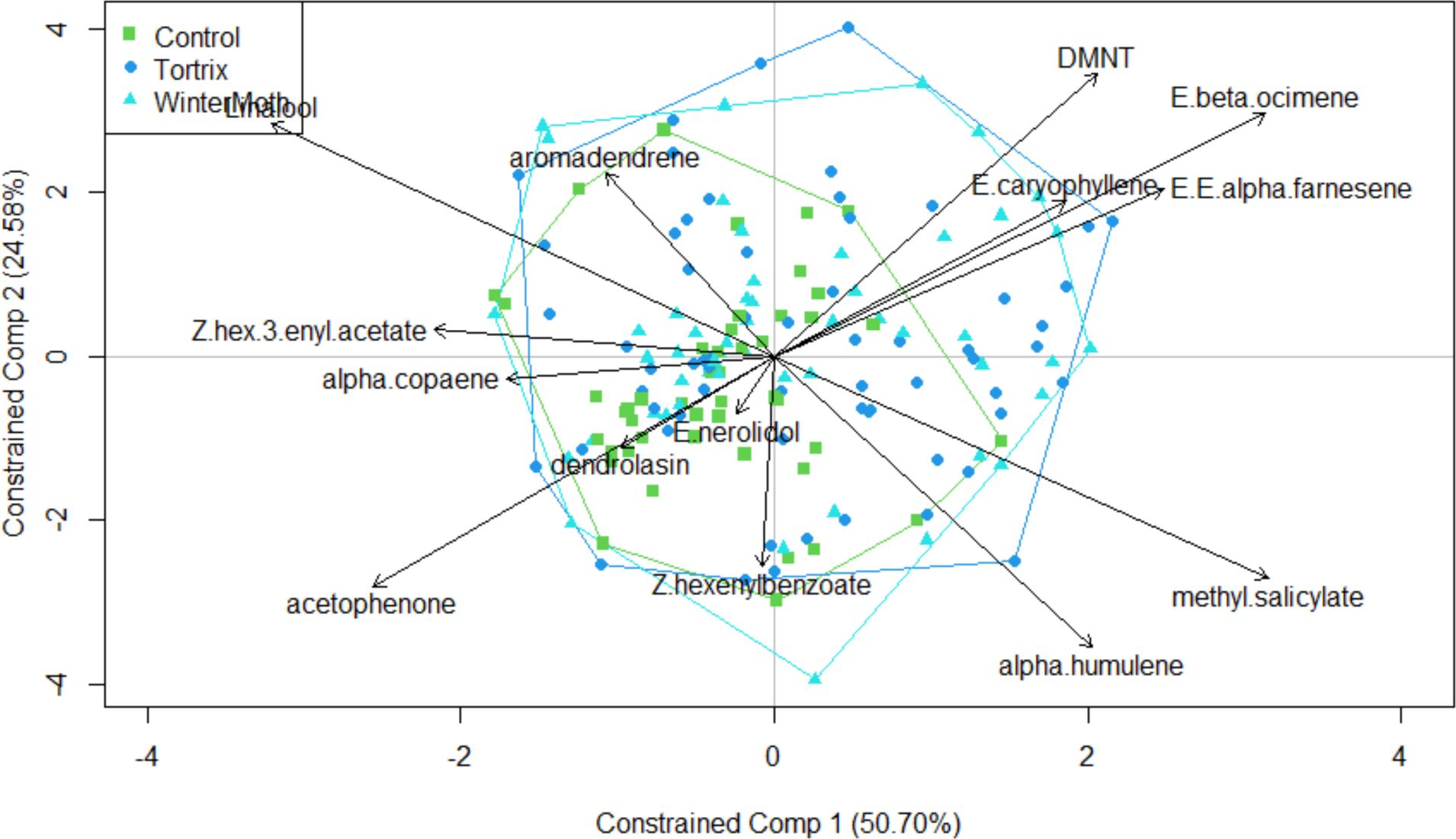
Biplot of VOC emissions shows predicted overlap in compounds between infested (green oak tortrix = blue, winter moth = light blue) and control trees (green), with infested trees extending beyond the area of the controls. VOCs extending clearly beyond the boundaries of the infested trees were considered as HIPVs. Aromadendrene and (*Z*)-hex-3-enyl benzoate were considered borderline significant, still being within the boundaries of the control trees, but near the border. The compounds dendrolasin and (*E*)-nerolidol were clearly within the boundary of the control trees and therefore were not considered as specific HIPVs in our study.

To visualize the patterns in HIPV emissions, the means (± SEM) for the compounds found to be significant and nearly significant in the previous analysis were graphed (figure 2). There was a general pattern for concentrations in the infested buds to increase compared to the control buds. The exception to this pattern was (*Z*)-hex-3-enyl acetate, which appeared to decrease over time (figure 2I).

**Figure 2.**
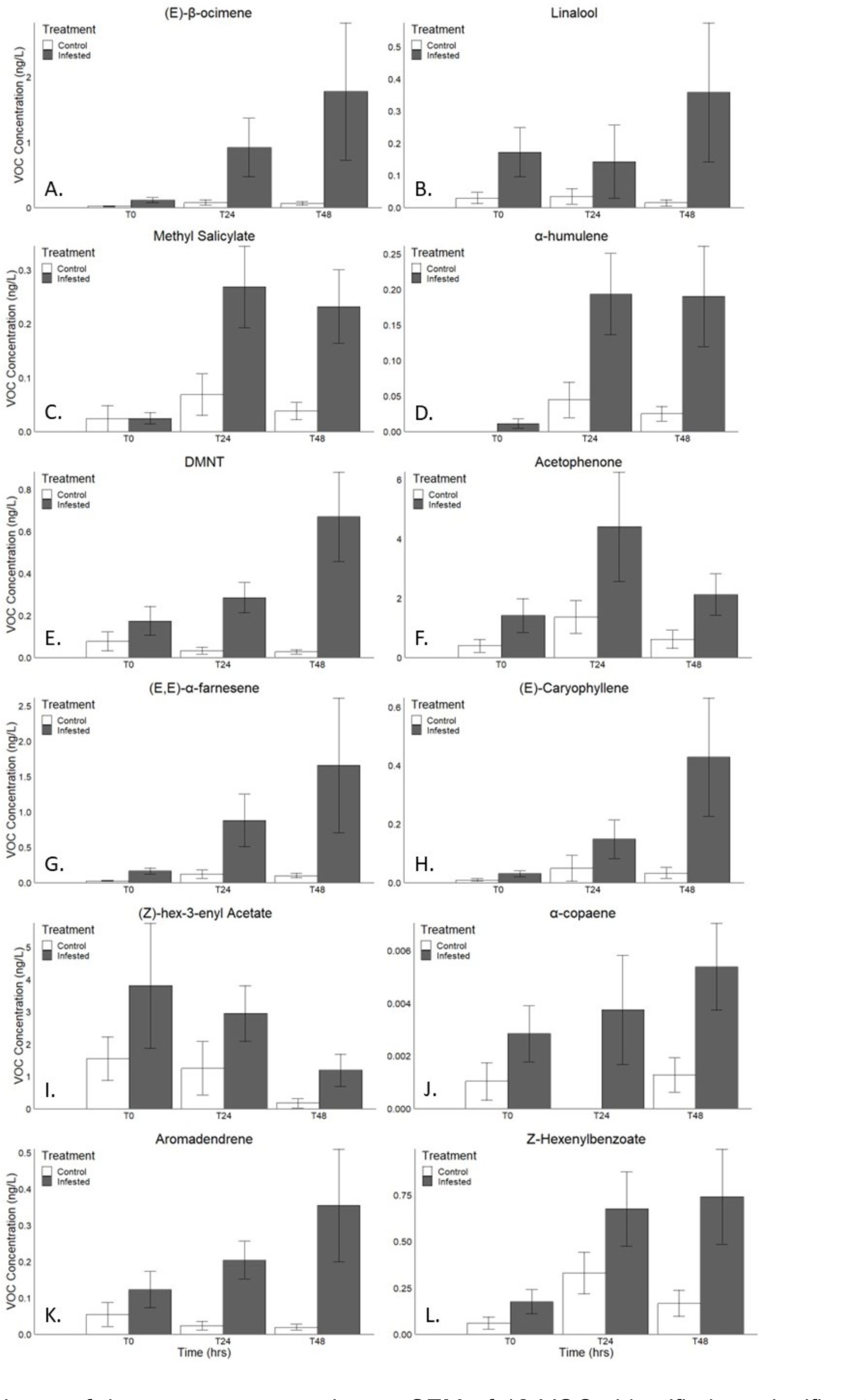
Comparisons of the mean concentrations ± SEM of 12 VOCs identified as significant or near significant from the analysis. While the interaction between time and treatment was not significant, a significant difference between time (*p* = 0.001) and between treatments (*p* = 0.014) was detected. Visualization of the data shows an increase over time within the infested trees (gray bars), with the exception of (Z)-hex-3-enyl acetate (panel I), and higher concentrations in the infested trees compared to the uninfested trees (white bars).

### 2018 Caterpillar Damages

After 96hr of infestation, green oak tortrix caused damage to an average of 23.58 ± 10.48 mm^2^ of leaf area while winter moth caterpillars caused damage to an average of 24.07 ± 9.55 mm^2^ corresponding to relative damages of 4.10 ± 2.02 and 3.83 ± 1.13 %, respectively. Relative damages ranged from 0.12 % to 14.6 %. There were no differences in the amount of leaf damages done by each species (*F*_1,19_ = 0.001, *p* = 0.97). When analyzing the effect of species and quantity of damage on the sum of HIPV emissions at 96h post-infestation, there is a significant, positive relationship with damaged leaf area (*F*_1,17_ = 8.62, *p* = 0.009, r = 0.54, figure 3). Species (*F*_1,17_ = 3.25, *p* = 0.09) and the interaction between species and damaged leaf area (*F*_1,17_ = 0.76, *p* = 0.39) were not significantly correlated with HIPV emissions.

**Figure 3.**
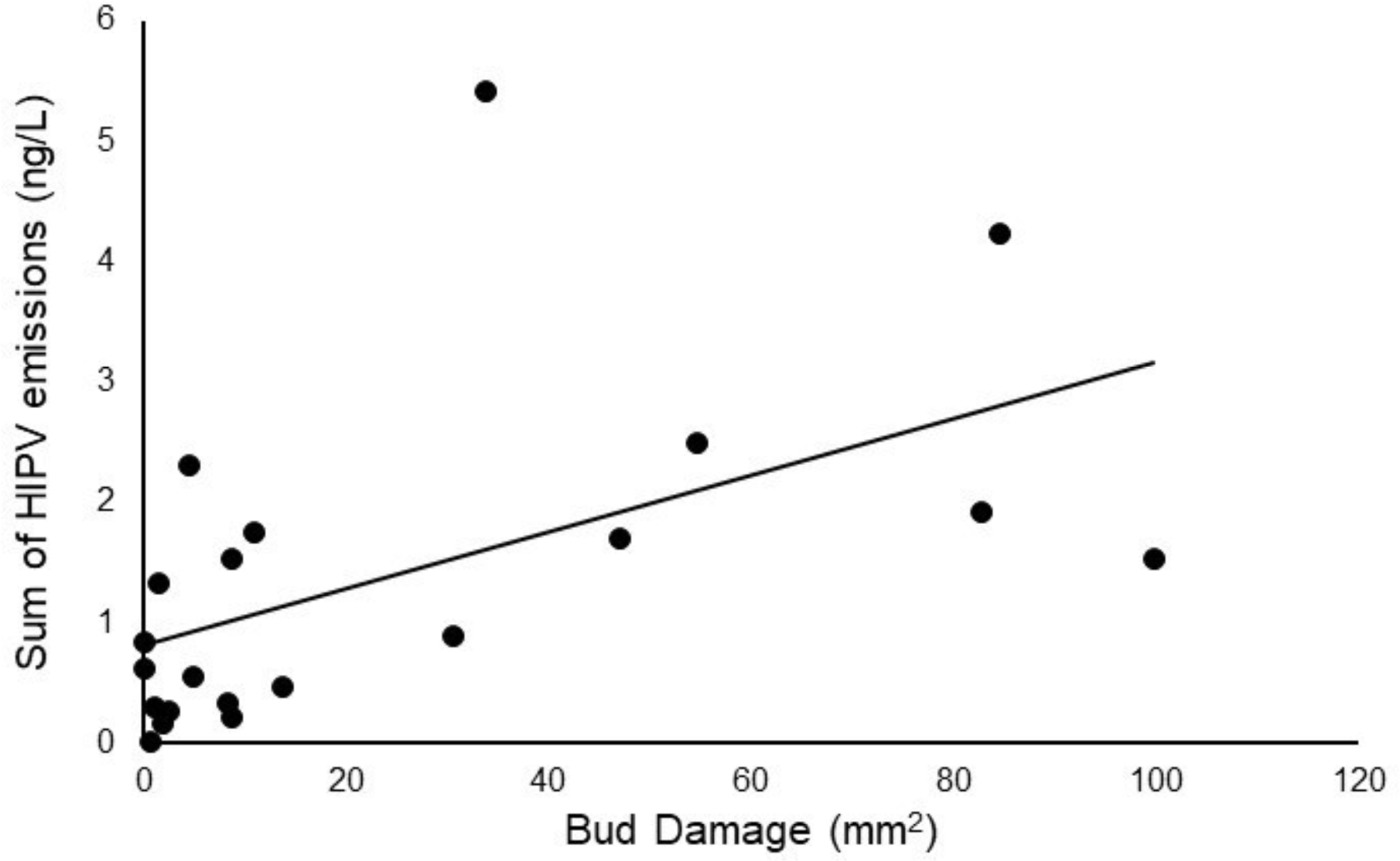
As the area of leaf damage increased, the sum of the VOC concentrations identified in 2018 samples also increased (r = 0.54). Species of caterpillar did not affect the quantity of damaged leaf area or the sum of HIPV emissions, suggesting the buds were responding to caterpillar herbivory in the same way, regardless of species.

## Discussion

Downy oak is known to be a strong constitutive emitter of isoprene (Rodríguez-Calcerrada *et al*. 2013), a compound we did not observe in our study on bud emissions. In fact, isoprene is produced only in mature leaves after induction by high temperatures (Monson *et al*. 1994) and therefore is irrelevant as a semiochemical in trophic relationships at earlier stages (see also for example Müller et al. 2015). Except for dendrolasin and (*E*)-nerolidol, all HIPVs released from Downy oak during bud break were significantly enhanced by caterpillar feeding, with a positive correlation between quantity of bud damage and HIPV production. This strongly suggests that these emissions were essentially non-constitutive. Indeed, DMNT, ocimenes, linalool, farnesenes, methyl-salicylate and hexenyl-acetate are among the most commonly observed HIPVs (for overviews, see e.g. McCormick *et al*. 2012; Dudareva *et al*. 2013; Turlings & Erb 2018). The minor emissions of these VOCs observed in control buds can be explained by some unavoidable mechanical stress they experienced during chamber installation and/or the presence of endo or epiphytical microorganisms (Junker & Tholl 2013). (*E*)-nerolidol is also a commonly reported HIPV. The non-significant link to caterpillar feeding of this VOC as well as of dendrolasin observed in the present study is likely due to their low volatility, making it difficult to measure emissions quantitatively. Both are oxygenated sesquiterpenes known to be chemically instable and to adhere and dissolve in surfaces including those of the plant tissues (Helmig *et al*. 2004; Niinemets *et al*. 2011). Moreover, (*E*)-nerolidol is the natural precursor of DMNT (Tholl *et al*. 2011) and therefore its emissions might evolve differently in response to biotic stresses. Generally, the induction kinetics of HIPV emissions can largely differ among individual compounds. Green leaf volatiles such as hexenyl-acetate and other volatile derivatives of fatty acid breakdown are almost instantaneously formed upon injury and rapidly disappear when stress ceases (see e.g. Staudt *et al*. 2010). By contrast, the induction of HIPV terpenoïds usually involves gene activation following a signaling cascade, which takes hours to days (Bonaventure *et al*. 2011; Joo *et al*. 2018). In line with this, our results show that most of the emitted terpenoïds steadily increased during 96 h enclosure period while hexenyl acetate emissions rather decreased. This decrease over time, which was also observed in the control buds, suggests that the hexenyl acetate emissions were induced in part already during the installation of the branches in the headspace and that many of the caterpillars reduced their feeding activities during the confinement, as indicated by their absence in some buds at the end of the experiment.

It has been hypothesized that specialist herbivores induce different defensive responses than generalist herbivores, because specialists gained the ability to tolerate direct plant defenses by sequestering phytotoxins (Ali & Agrawal 2012; McCormick *et al*. 2012; Rowen & Kaplan 2016). They may also manipulate the host metabolism to their benefit (Lin *et al*. 2022) and even use HIPVs as cues to choose the right host plant. For example *Tortrix viridana* females are more attracted by the VOC profiles emitted by susceptible pedunculate oak ecotypes than by the profile of resistant ecotypes, whose leaves contain secondary metabolites reducing larval growth and survival (Ghirardo *et al*. 2012; Bertić *et al*. 2021). Owing to their adaptation, specialists are expected to inflict more feeding damage on host trees than generalists, which in turn could induce more HIPVs, possibly favoring the plant’s indirect defenses against the herbivore. However, meta analyses on literature data provide only limited support for this hypothesis, and revealed that comparative studies sometimes confounded specialization (diet breadth) and feeding guild (i.e., chewing insects vs sap feeders; (Ali & Agrawal 2012; Rowen & Kaplan 2016). Sap feeders such as aphids indeed induce less and different HIPV patterns than chewing insects, because they inflict little to no tissue damage but are potential vectors of pathogens (e.g., Staudt *et al*. 2010). Here, we found no detectable differences in the VOCs produced in response to herbivory by the generalist *Operophtera brumata* and the oak specialist *Tortrix viridana*. HIPV production was correlated with damage, but it was similar for both caterpillar species. This suggests that the induction of HIPVs depended predominantly on damage derived factors rather than on insect specific elicitors (Arce *et al*. 2021). Indeed, Mithöfer *et al*. (2005) showed that mechanical wounding simulating caterpillar foraging induces VOC emission patterns similar to real herbivory.

One result we consider having a significant ecological relevance is that total HIPV concentrations were related to the amount of damages inflicted to the buds. Positive correlations between biotic, stress-induced VOC emission and the degree of damage has been previously reported (Niinemets *et al*. 2013). Since alarm signals emitted by plants are considered cry-for-help (Dicke 2009), releasing more HIPVs when plants are more severely attacked (Girling et al 2011), may seem adaptive as studies have shown that parasitic wasps were more attracted to plants emitting more HIPVs (Guerrieri et al. 1999). Most studies were however conducted with plants that were already in full leaf, and which were attacked by mature insects (but see for example Hare 2010). In the present study, we show that emissions proportional to infestation already occur at a much earlier stage, when oaks are still in their bud stages and are attacked by newly hatched caterpillars. Those caterpillars are only about 1 mm long, hide inside buds (DuMerle & Mazet 1983), and are still too small to be considered preys for predators like birds that need large biomass of insects to feed their chicks (Blondel et al. 1999). Furthermore, predation of caterpillars by birds at this early stage would likely entail the complete loss of the infested buds (Betts 1955), more so than a caterpillar attack by specific parasitoid wasps. There is thus a potential conflict between plants that need their precious buds to be protected from irreversible damages, and insect predators that may not be very interested by newly hatched caterpillars, or cause even more severe damages than caterpillars. From the plants’ perspective, the net benefits of HIPV emission from buds via tri-trophic interactions thus remains to be clarified (Dicke & Baldwin 2010). One potential explanation comes from recent studies suggesting that those early emissions already elicit physiological and behavioural reactions from insectivorous songbirds that may use these odours as signals to modulate their reproductive timing and effort (Graham et al. 2021, Caro et al. 2023). Birds could for example modulate their clutch size, i.e. the number of chicks they intend to raise, based on the amount of HIPVs they detect at this early stage. In this scenario, bud HIPVs would be used as a cue to infer the future available biomass of caterpillars. Therefore, even if the immediate benefits for plants of emitting HIPVs while they are still in buds is not clear yet, they might at least guarantee that there will be enough manpower around to rid them of their parasites at a later stage.

In the present study, we did not investigate whether HIPVs were only emitted from the damaged buds or also from other non-attacked plant parts. Many studies have shown that HIPVs can be systemically emitted upon stress (e.g. Röse & Tumlinson 2004) possibly increasing its signaling range for receivers (McCormick *et al*. 2012). However, HIPVs released into the air masses are subject to rapid turbulent transport, mixing, and chemical reactions with other trace gases (Conchou *et al*. 2019). Therefore, the distances odor plumes can travel and be perceived by other organisms are highly variable, independent of the shape and the strength of the emission source (Douma *et al*. 2019; Pannunzi & Nowotny 2019). Furthermore, few studies have quantified local and systemic emission rates. In a study of holm oaks infested with gypsy moth, Staudt & Lhoutellier (2007) reported that HIPV emission rates from uninfested leaves were about an order of magnitude lower than rates from infested leaves, indicating that the source of HIPVs is essentially, although not exclusively, local. From the perspective of a caterpillar predator using HIVPs for foraging, locally emitted HIPVs should be much more efficient for spotting their prey in the close range (Volf *et al*. 2021), unless cues other than olfactory ones play a role.

In conclusion, few studies exist that measure the HIPVs of tree buds in response to herbivore attacks. Our study suggests that, even when herbivorous insects are too small to be considered food or host for insect predators, Downy oak are upregulating the production of HIPVs. The purpose of this upregulation remains unclear. Thus, future studies should investigate whether and how herbivorous insect predators use these cues, and whether there are reciprocal benefits to the tree itself.

## Statements and Declarations

We thank Marjorie Simean and Selim Ben Chehida for their help during the experiments; Christophe de Franceschi and Bart van Lith for providing the caterpillar eggs, and people from the TE (Terrains d’Expériences) platform at CEFE for their logistic help with maintaining the trees. This work was funded by a grant from the Agence Nationale de la Recherche awarded to SPC (grant number ANR-15-CE02-0005-01). The authors have no competing interests to disclose.

